# High contrast probe cleavage detection on porous silicon biosensors via quantum dot labeled DNA probes

**DOI:** 10.1101/2020.10.07.330589

**Authors:** Rabeb Layouni, Michael Dubrovsky, Mengdi Bao, Haejun Chung, Ke Du, Svetlana V. Boriskina, Sharon M. Weiss, Diedrik Vermeulen

## Abstract

Using porous silicon (PSi) interferometer sensors, we show the first experimental implementation of the high contrast probe cleavage detection (HCPCD) mechanism. HCPCD makes use of dramatic optical signal amplification caused by cleavage of high-contrast nanoparticle labels on probes instead of the capture of low-index biological molecules. An approximately 2 nm reflectance peak shift was detected after cleavage of DNA-quantum dot probes from the PSi surface via exposure to a 12.5 nM DNase enzyme solution for 2 hrs. This signal change is 20 times greater than the resolution of the spectrometer used for the interferometric measurements, and the interferometric measurements agree with the interferometric response predicted by simulations and fluorescence measurements. These proof of principle experiments show a clear path to real-time, highly sensitive and inexpensive point-of-care readout for a broad range of biological diagnostic assays that generate signal via nucleic acid cleavage.

## 1. Introduction

Breakthroughs in nucleic acid testing are needed to meet today’s demand for SARS-CoV-2 tests and to ensure resilience to future pandemics. Commercially-available assays for pathogen nucleic acid diagnostics heavily rely on biological amplification steps to increase the amount of target material via replicating ribonucleic acid (RNA) or deoxyribonucleic acid (DNA) of the pathogen genome [1]–[3]. Biological amplification steps increase the test complexity and processing time and may sometimes generate false positive results [4]. On the other hand, adaptation of conventional photonic and plasmonic biosensing platforms [5]–[7] for early-stage detection poses challenges of returning false negative results if the amount of the viral RNA in the sample is below the sensor detection limit. Standard photonic biosensors make use of surface functionalization by probes that can selectively capture target molecules, and the detection limit is defined by how many molecules should be captured before a response from the sensor could easily be distinguished from its inherent noise. Furthermore, when new pathogens emerge, it may take up to 6-12 months to develop new probes specific to the target [8].

We have previously demonstrated via simulations that high contrast probe cleavage detection (HCPCD) can provide a highly sensitive readout based on removing probes from the photonic biosensor surface [9]. The HCPCD sensing modality is the reverse of standard binding/adsorption-based label-free optical biosensing methods [8], which depend on the small difference in refractive index between samples such as blood or saliva and biomolecules binding to the sensor surface. HCPCD offers all the advantages of miniaturization and optical signal amplification of photonic sensors, while significantly increasing the signal-to-noise ratio if the probes are chosen such their removal generates a much stronger signal that the attachment of the same amount of low-index biomolecular targets. Semiconductor nanocrystals, quantum dots, or metal nanoparticles can serve as high-contrast probes, which can be attached to the sensor surface via known techniques, thus eliminating the lengthy phase of the new probe chemistry development. Furthermore, since sensor functionalization with the nanoprobes attached to RNA/DNA tethers can be done prior to the sample collection from the patient or the environment, the time to detection can also be shortened significantly. During the testing process, the DNA/RNA tethers can be cleaved by introduction of enzymes capable of hydrolyzing phosphodiester bonds that link nucleotides in DNA/RNA chains, such as deoxyribonuclease (DNase) [10] or Clustered Regularly Interspaced Short Palindromic Repeats-associated complexes (CRISPR-Cas) [3], [11].

In order to validate the working principles of HCPCD and its possible applications in RNA and DNA detection, DNase enzyme sensing experiments were performed using a porous silicon (PSi) interferometric sensor functionalized with either high-contrast ds-DNA-quantum dot (Qdot) or low-index ds-DNA-fluorescein (FAM) probes. PSi interferometer sensors are a well-studied promising simple platform for low-cost diagnostics [9], [12]–[14]. This platform offers large surface-to-volume ratio, simple optical reflectance readout, and surface functionalization by using well-developed surface chemistries. Although DNA detection has been demonstrated using PSi sensors, challenges in achieving high sensitivity detection remain. Typically, experimental detection of µM concentrations of DNA are reported. While signal amplification through quantum dot attachment to target DNA [15], [16] and electrokinetic focusing [17] to concentrate target DNA lead to improved sensitivity, these approaches are not compatible with simple, real-time diagnostics. In this work, we not only demonstrate that HCPCD detection is a promising approach to improve the sensitivity of PSi sensors in a straightforward manner, but we also importantly establish the feasibility of the HCPCD detection approach for future implementation on resonant silicon-based optical sensors, which have the potential to reach the sensitivity necessary for amplification-free PCR-quality assays [9].

## 2. Working Principle of PSi Reflective Interferometer with HCPCD Mechanism

Single layer PSi interferometers work on the principle of thin film interference. Light reflected from the top (air/PSi) and bottom (PSi/silicon substrate) interfaces of the PSi layer interfere, giving rise to characteristic Fabry-Perot reflectance fringes (Fig. 1(a)). The fringe positions and the periodicity of the interferometric pattern are determined by the effective optical thickness of the PSi film:

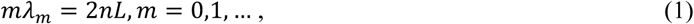

where *n* is the effective refractive index of the PSi layer, *L* is the layer thickness, and λ_*m*_ is the spectral position of the *m*-th fringe maximum. Since the average pore diameter is much smaller than the wavelength of light, an effective medium approximation can be used to estimate the effective refractive index of the PSi film.

**Fig. 1.**
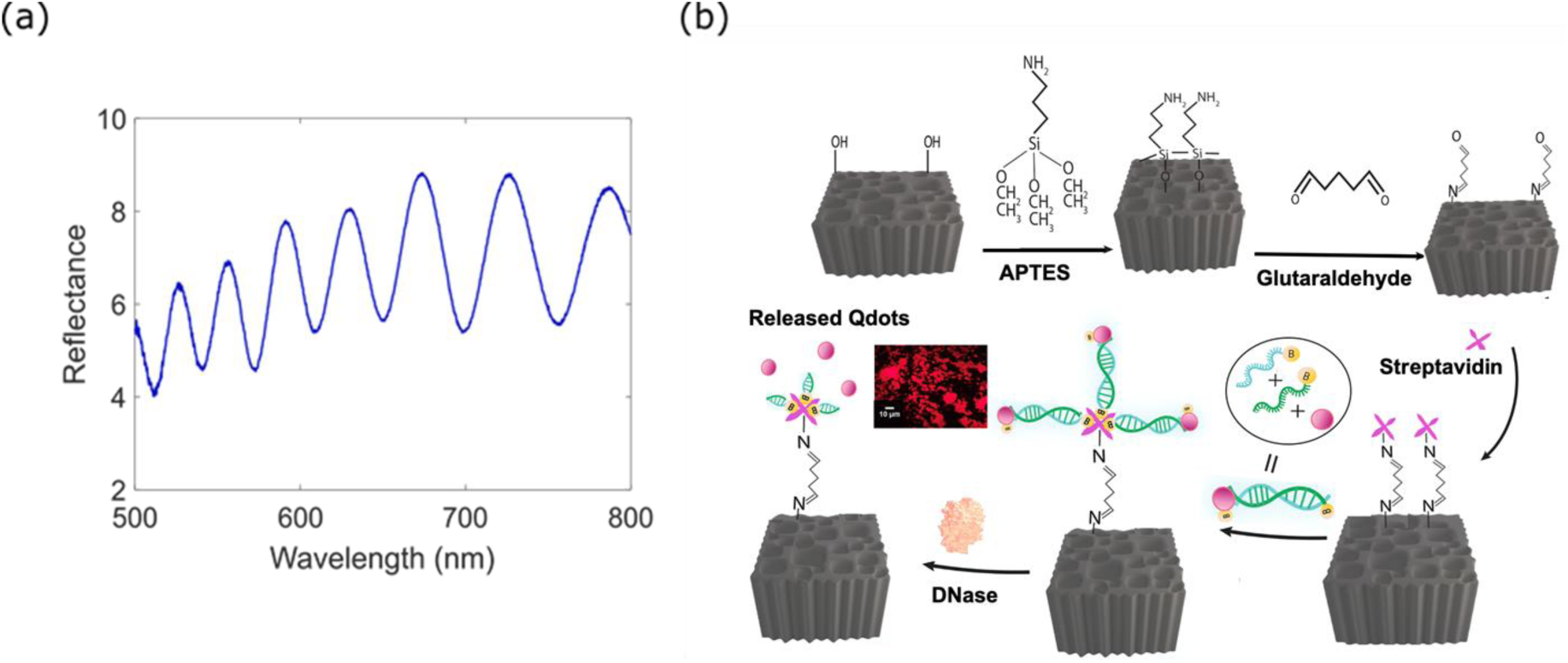
(a) Interference spectrum collected from an oxidized PSi layer under normal illumination by a broadband white light source. (b) Schematic of the HCPCD-based reflectance interferometry detection platform with a PSi layer as the optical transducer. The PSi layer is functionalized with quantum dot-labeled probes using the appropriate surface chemistry and the biotin-streptavidin conjugation mechanism, as described in the methods section. The Qdot probes are then cleaved by the exposure of the sensor to the DNase enzyme and washed away, which generates a measurable optical signal change in the form of the Fabry-Perot fringes spectral shift. The inset shows a fluorescence image of the Qdots.

At the same time, the pore size is chosen to be larger than both the molecular target and the molecules used in the surface functionalization process, to allow sufficient space for the molecular species to infiltrate into the large surface area of the porous material. When molecular species enter the pores, the effective refractive index of the PSi layer increases, and the fringes redshift. Conversely, when species (or dielectric nanoparticle probes) are removed from the pores, the effective refractive index of PSi decreases and the fringes blueshift. In the HCPCD detection approach, as illustrated in Fig. 1(b), a PSi interferometer is first functionalized with high-index biotin-dsDNA-Qdot probes, leading to a redshift of the reflectance fringes. When an enzyme is used to cleave the DNA, leading to removal of the Qdots, the reflectance fringes blueshift. The magnitude of the reflectance shifts are proportional to the quantity of material being added to or removed from the pores, and can provide a mechanism for a sensitive and quantitative optical readout. In the following sections, we describe the methodology in detail.

## 3. Experimental Methods and Procedure

### 3.1 Porous silicon fabrication and surface modification

PSi thin films used for the sensing experiments in this work were fabricated by electrochemical etching in a solution of 15% hydrofluoric acid in ethanol at a current density of 70 mA/cm^2^ for 100 s. We note that prior to fabricating these PSi films, a sacrificial PSi layer was first etched and then removed in NaOH solution to ensure that there were large pores openings at the surface [18]. After the removal of the sacrificial layer and again after formation of the PSi film used for sensing, chips were washed with ethanol and dried with nitrogen. Surface modification followed a previously established procedure [19]. First, to provide surface passivation and prepare for silanization, freshly etched PSi chips were thermally oxidized at 800 °C for 30 min. Chips were subsequently rinsed with ethanol and dried with nitrogen. The thickness and porosity of the PSi layer were measured by scanning electron microscopy (SEM), and found to be 3.99 mm and 73%, respectively.

The oxidized samples were incubated in freshly prepared 4% 3-aminopropyltriethoxysilane (APTES) solution (1 mL deionized water, 920 μL methanol and 80 μL APTES) for 15 min. The samples were then soaked in methanol for 15 min. Excess APTES was removed by rinsing with deionized (DI) water and ethanol. The samples were dried with nitrogen, and then annealed at 150 °C for 10 min, rinsed with methanol and dried with nitrogen. Next, the samples were incubated in a 2.5% glutaraldehyde solution (50 μL of 25% glutaraldehyde and 950 μL HEPES buffer, pH =7.4) for 30 min. Excess glutaraldehyde was removed by incubating each sample in 1 μL of sodium cyanoborohydride in 100 μL HEPES buffer solution for 30 min, which is also necessary to stabilize the Schiff base. Finally, streptavidin was attached to the surface by exposing the samples to a 10 μM streptavidin solution in PBS buffer (pH=7.4) for 30 min, and the Schiff base was stabilized by repeating the reducing step with sodium cyanoborohydride. To minimize non-specific binding, the sample was incubated in 3 M ethanolamine in buffer with pH=9.0 for 30 min. The sample was then thoroughly rinsed with DI water and dried with nitrogen.

### 3.2 Synthesis of quantum dot and FAM dye labeled probes

To synthesize biotin-dsDNA-Qdot probes, 0.2 µM Qdots (QdotTM 605 ITKTM Streptavidin Conjugate Kit, ThermoFisher Scientific) were first mixed with 5 µL of 100 µM biotinylated ssDNA (5’ biotin-CTGTCGTGGTTCTAGGGAGTCAAGAAGGAGCAGTTCACATTT 3’; IDT, Inc.) to reach a total volume of 36.2 µL. The mixture was incubated in the dark at room temperature for 15 min. Next, 10 µL of complementary biotinylated ssDNA (IDT, Inc.) with a concentration of 100 µM was added and reacted with the mixture under mild shaking with a tube rotator (BS-RTMI-1, STELLAR SCIENTIFIC) for 90 min.

To synthesize FAM dye labeled probes, two biotinylated complementary DNA strands at 100 µM concentration with FAM dye tags (IDT, Inc.) were hybridized via a 90-minute incubation under mild shaking with the tube rotator.

### 3.3 Probe attachment to porous silicon

Biotinylated dsDNA-Qdot probes were attached to the streptavidin-modified PSi surface by incubating the sample in 24 µL of 135 nM probe solution overnight. The footprint of PSi exposed to the probe solution was 0.25 cm^2^. After incubation, the supernatant was collected, and the sample was rinsed thoroughly with DI water and dried with nitrogen. The same procedure was repeated on a separate PSi sample using 135 nM biotinylated dsDNA-FAM probes.

### 3.4 DNase cleavage of the probes

Each PSi sample was incubated in DNAse I solution (2 µL DNAse solution (1U/µL), 2 µL buffer and 16 µL DI water) for 2 h at 310 K in a moist environment. The samples were rinsed thoroughly with DI water and dried with nitrogen.

### 3.5 Measurements

Reflectance measurements were carried out using a Newport Q series lamp and fiber-coupled Ocean Optics USB-4000 spectrometer. Attenuated total reflectance Fourier transform infrared (ATR-FTIR) spectroscopy measurements were taken using a Hyperion 2000/Vertex 70v system with a silicon prism. Fluorescence measurements were performed on a Jobin Yvon Fluorolog-3 FL3-111 spectrofluorometer.

### 3.6 Simulations

We validated the experimental measurements with transfer matrix calculations [20]–[22]. The refractive index of the functionalized PSi matrix with and without probes was modeled with the two-component and the three-component Bruggeman’s effective medium theory [20], [23], [24], respectively. The thickness of the organic layer that is used to capture the Qdot probes (APTES-glutaraldehyde-streptavidin) was estimated to be 3.5 nm based on an assumed 50% surface coverage of streptavidin molecules in the pores and near monolayers of APTES and glutaraldehyde. This leaves a 54% volume fraction of air in the total sensor volume, assuming the initial PSi porosity of approximately 73%. According to Bruggeman’s effective medium theory, the effective refractive index of the PSi matrix with the organic layer (*n*_*m*_) can be calculated as follows:

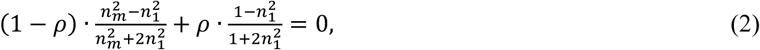

where *ρ* = 0.54 is the porosity of the functionalized PSi layer, the void space refractive index is taken as 1 for air, and *n*_1_ is Bruggerman-model effective refractive index of this layer, which can be found from the fast Fourier Transform (FFT) of the experimental spectrum [14] taken prior to attachment of the biotin-dsDNA-Qdot probes, as shown in Fig. 2(a).

**Fig. 2.**
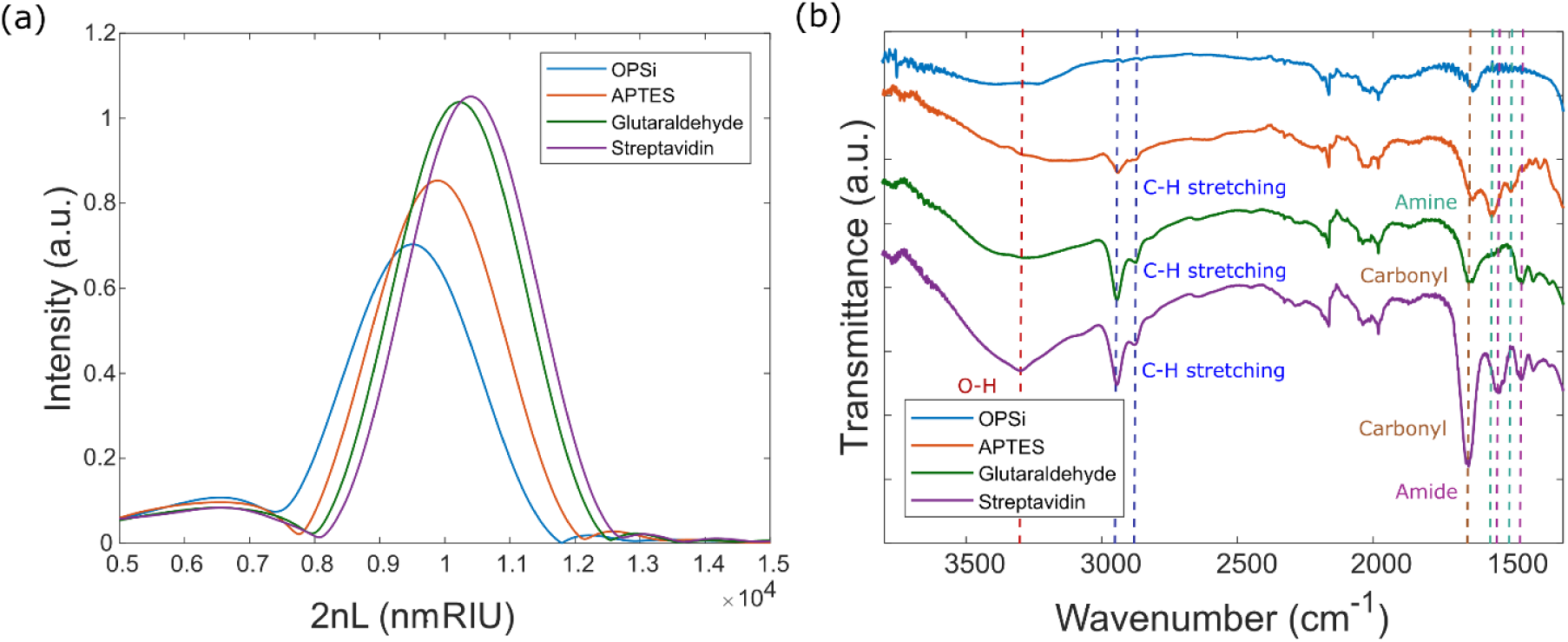
(a) FFT of reflectance spectrum of PSi after various surface modification steps. (b) Corresponding ATR-FTIR spectra of PSi measured after each surface modification step in (a); important peaks are labeled.

The addition of biotin-dsDNA-Qdot probes – which can only occupy the space within the air pores – was modeled via the three-component Bruggeman’s model:

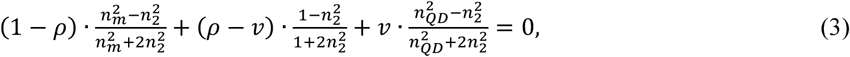

where *v* is the unknown volume fraction occupied by the biotin-dsDNA-Qdot probes, *n*_*QD*_ is the effective index of the composite biotin-dsDNA-Qdot probe, and *n*_2_ is Bruggeman-model effective refractive index of the functionalized porous layer with the probes, which can be found from the FFT of experimental spectrum taken after attachment of the probes. The effective refractive index of the biotin-dsDNA-Qdot probes, *n*_*QD*_ = 1.52, was found by applying the two-component Bruggeman model (Eq. 2) to simulate a probe composed of a Qdot with a refractive index of 2.5 [25] surrounded by the surface coating of biological material with a refractive index of 1.45 [26]. The Qdot is modeled as an 8 nm radius sphere [27]. The volume of the biological material associated with each Qdot was found by combining the volume of 7 streptavidin molecules [28], each modeled as a 5 nm radius sphere, and 100 biotinylated dsDNAs with 21 base pairs, each modeled as having a volume of 2.8×10^−8^ µm^3.^ It is assumed that the dsDNAs not only quench the binding sites on the Qdot-streptavidin complexes, but also that the excess dsDNA in the probe solution (see section 3.2) is available to directly bind to the surface of the streptavidin-functionalized PSi.

Combining all these effective index approximations, the filling fraction of the probes attached in the void space of PSi was calculated based on the comparison to experimental spectra taken after the Qdot attachment step. The simulated probe filling fraction was adjusted such that the simulated spectrum after probe attachment matched the experimentally measured spectrum. The total number of Qdots bound inside the PSi was then calculated and compared to the total number of Qdots estimated to be in the pores based on the change of the measured fluorescence intensity of the probe solution before and after exposure to PSi, as discussed in section 4.

## 4. Results and Discussion

The as-anodized PSi films used in this work had a thickness of approximately 3.99 µm, porosity of 73%, and average pore diameter of approximately 50 nm. To prepare the surface for probe attachment, the films were first thermally oxidized and then APTES, glutaraldehyde, and streptavidin were sequentially attached to the oxidized PSi (OPSi) surface. Reflectance measurements and ATR-FTIR measurements were taken after each surface modification step, as shown in Fig. 2(a,b). Additionally, after the streptavidin surface modification step, the PSi was soaked in DI water for 24 h and then remeasured to ensure that the surface modification was stable before proceeding to probe functionalization.

The reflectance spectra of the PSi thin films are characterized by Fabry-Perot fringes (see, for example, Fig. 1(a)). The spacing of these fringes is determined by the optical thickness of the PSi film. For large changes in the optical thickness of the PSi film that cause large fringe shifts (e.g., due to the attachment of a large number of molecules in the pores), it can be challenging to analyze the exact spectral change due to the possibility of the fringes shifting by more than the fringe spacing. Accordingly, for the initial surface modification steps, we analyzed the optical thickness changes by using the reflective interferometric Fourier transform spectroscopy technique [14]. In this approach, taking the FFT of the reflectance spectrum gives a peak whose position corresponds to twice the optical thickness of the PSi film (i.e., 2*nL* where *n* is the effective refractive index of the PSi and *L* is the physical thickness of the PSi film). As shown in Fig. 2(a), the 2*nL* peak position increases after each surface modification step, suggesting that APTES, glutaraldehyde, and streptavidin are attached to the OPSi. Negligible shift in the 2*nL* peak position was measured after the DI water soak following streptavidin attachment, suggesting that the PSi surface modification is stable.

ATR-FTIR measurements confirm that the measured optical thickness changes are indeed caused by the different molecular attachments. N-H bending peaks around 1580 cm^−1^ and 1650 cm^−1^, shown in Fig. 2(b), indicate the presence of amine in APTES. An increase in C-H stretching peaks at 2863 cm^−1^ and 2966 cm^−1^ and appearance of a carbonyl peak at 1715 cm^−1^ confirm the presence of glutaraldehyde on the surface. Finally, the attachment of streptavidin is shown by O-H, carbonyl, and amide bands peaks at 3300 cm^−1^, 1721 cm^−1^, and 1470 cm^−1^ and 1600 cm^−1^, respectively.

After incubating the streptavidin-modified PSi in the 135 nM biotin-dsDNA-Qdots probe solution, the supernatant was collected for fluorescence measurements and the reflectance of the PSi was measured after the film was rinsed and dried. As shown in Fig. 3(a), the Fabry-Perot fringes redshift by approximately 3.1 nm after exposure to the probes, suggesting the biotin-dsDNA-Qdots were attached to the streptavidin-modified PSi. Note that due to the excess biotinylated DNA in the probe solution, some of the streptavidin molecules on the PSi surface may capture biotinylated DNA instead of biotin-dsDNA-Qdots. Fluorescence measurements comparing an equal volume of the supernatant with 135 nM biotin-dsDNA-Qdots solution that was not exposed to PSi are shown in Fig. 3(b). A clear decrease in fluorescence intensity for the supernatant confirms that the biotin-dsDNA-Qdots were attached to the streptavidin-modified PSi. To ensure the probes were not non-specifically trapped in the pores, a streptavidin-modified PSi film was exposed to 135 nM streptavidin-coated Qdots. As shown in Fig. 3(c,d), there was negligible change in reflectance after exposure to the streptavidin-coated Qdots and the fluorescence intensity of the supernatant of the streptavidin-coated Qdots exposed to the PSi was the same as that of an equal volume of 135 nM streptavidin-coated Qdots solution not exposed to PSi. Hence, we conclude that there is negligible non-specific probe binding occurring in the PSi film.

**Fig. 3.**
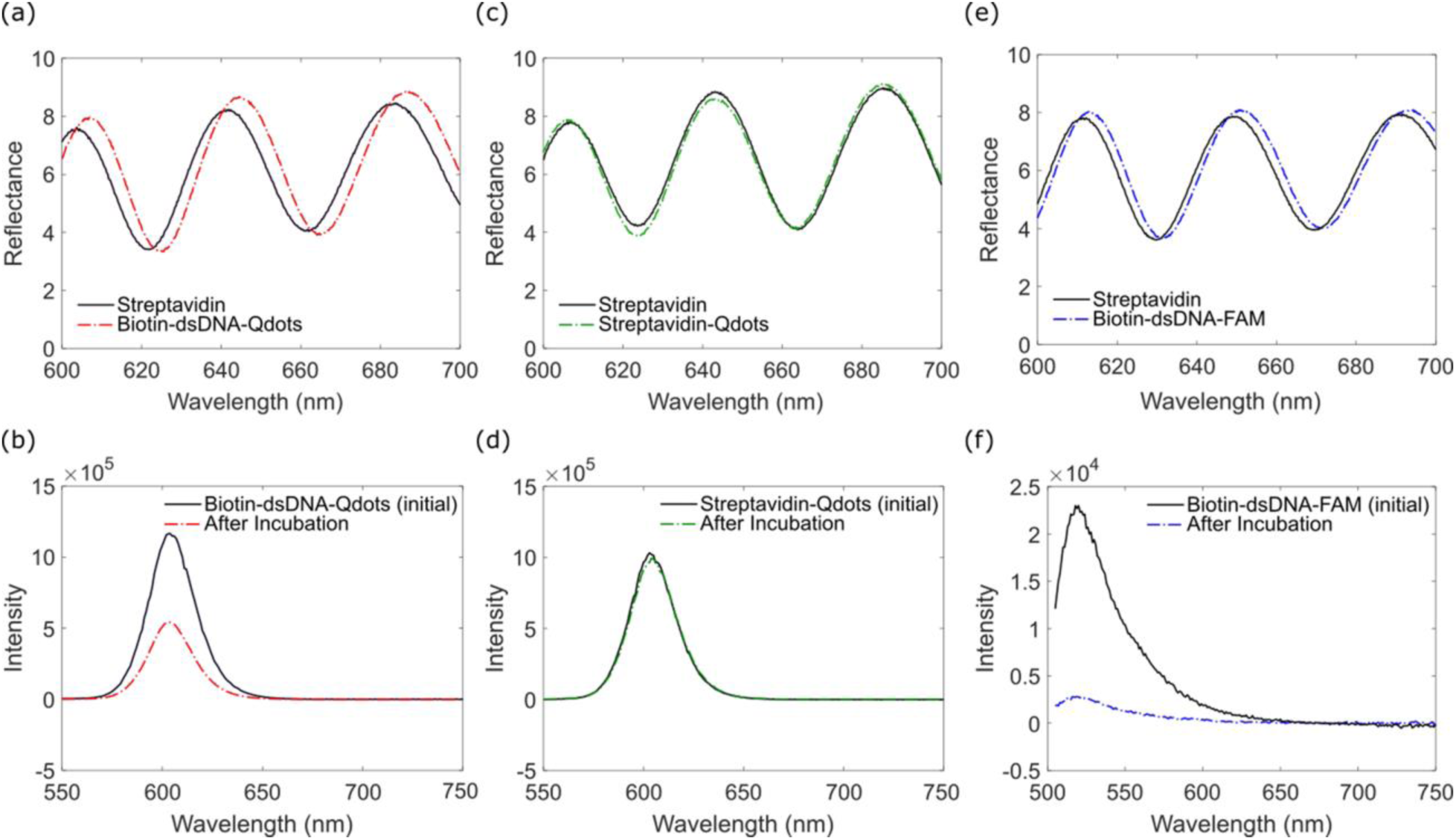
(a) Reflectance spectra before and after attachment of 135 nM biotin-dsDNA-Qdot probes to streptavidin-modified PSi. (b) Fluorescence intensity of 135 nM biotin-dsDNA-Qdots probe solution compared to equal volume of biotin-dsDNA-Qdots supernatant collected after exposure of the probes to streptavidin-modified PSi. The lower fluorescence intensity of the supernatant is consistent with probes being attached to the streptavidin-modified PSi during incubation. (c) Reflectance spectra before and after attachment of control solution of 135 nM streptavidin-coated Qdots to streptavidin-modified PSi. No change in reflectance suggests that there is no non-specific binding. (d) Fluorescence intensity of 135 nM streptavidin-Qdots solution compared to equal volume of streptavidin-Qdots supernatant collected after exposure to streptavidin-modified PSi. No change in fluorescence intensity is consistent with negligible non-specific binding of the Qdots in the PSi. (e) Reflectance spectra before and after attachment of 135 nM biotin-dsDNA-FAM probes to streptavidin-modified PSi. (b) Fluorescence intensity of 135 nM biotin-dsDNA-FAM probe solution compared to equal volume of biotin-dsDNA-FAM supernatant collected after exposure of the probes to streptavidin-modified PSi.

Figure 3(e) shows the reflectance measurements of a streptavidin-modified PSi film before and after attachment of 135 nM biotin-dsDNA-FAM probe solution: a 1.7 nm redshift results from attachment of these probes. Fluorescence measurements of the supernatant of the biotin-dsDNA-FAM probe solution after incubation on the PSi film showed decreased intensity compared to an equal volume of solution of the biotin-dsDNA-FAM probes that was not exposed to PSi (Fig. 3(f)). This decrease in fluorescence intensity is consistent with probe attachment to the streptavidin-modified PSi. Comparing the Fabry-Perot fringe shifts in Fig. 3(a,e), the redshift for probe attachment is nearly 2 times larger for the biotin-dsDNA-Qdots than the biotin-dsDNA-FAM, suggesting that a larger change in the optical thickness occurs when the biotin-dsDNA-Qdots are attached. This result indicates that the higher refractive index and larger size of the Qdots compared to FAM make them a superior high contrast label for the probes. This conclusion is confirmed in the DNase cleavage detection experiment, as discussed below.

As a proof of principle demonstration of HCPCD, DNase was used to cleave the dsDNA and release the probes from the pores. First, the experiment was carried out with the PSi sample functionalized with biotin-dsDNA-Qdot probes. Figure 4(a) shows a nearly 2.2 nm blueshift of the Fabry-Perot fringes that results after the Qdots and some portion of the DNA were removed from the PSi surface and rinsed out of the pores. The fringes do not shift fully back to their position prior to probe attachment as some portion of the biotinylated DNA remains attached to the streptavidin-modified PSi surface. Moreover, the DNase enzyme likely did not cleave every DNA tether and thus some probes may remain bound to the pore walls. The simulations discussed below and shown in Fig. 4(b) provide insights on the quantity of probes attached and subsequently cleaved from the PSi sample. Importantly, in the control experiment shown in Fig. 4(c), in which a biotin-dsDNA-Qdot-functionalized PSi film was soaked in DI water for 2 h – a duration equivalent to the DNase exposure – there was only a minor change to the reflectance spectrum (∼ 0.4 nm blueshift), which may be due to the fact that DI water and not specifically DNase-free water was utilized for the soak. Thus, we believe that the probe-functionalized PSi surface is stable and DNase leads to removal of material from the pores. When PSi functionalized with biotin-dsDNA-FAM probes was exposed to DNase solution, a nearly 1.2 nm blueshift of the Fabry-Perot fringes was measured, as shown in Fig. 4(d). The magnitude of the blueshift was much less for the PSi with biotin-dsDNA-FAM probes after exposure to DNase compared to that for the PSi with biotin-dsDNA-Qdot probes, confirming that Qdots are the preferable probe label compared to FAM dye for HCPCD sensing.

**Fig. 4.**
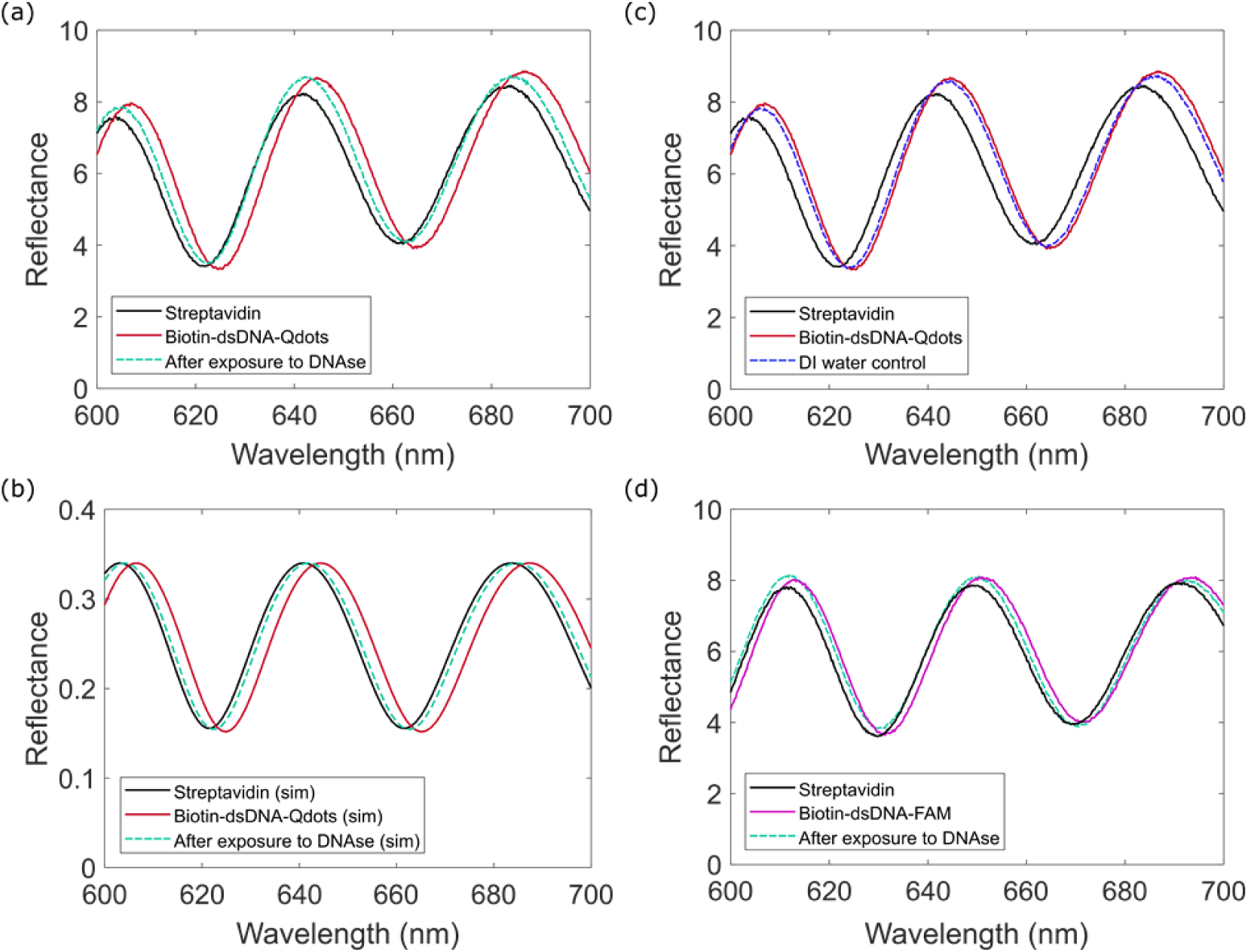
(a) Reflectance spectra showing redshift of Fabry-Perot fringes after biotin-dsDNA-Qdot probe attachment to streptavidin-modified PSi and blueshift of fringes after exposure to DNase and removal of DNA and Qdots from the PSi. (b) Simulated reflectance spectra showing redshift of Fabry-Perot fringes after 0.6 picomoles of biotin-dsDNA-Qdot probes are attached to streptavidin-modified PSi and blueshift of fringes after removal of approximately 67% of DNA-Qdots from the PSi. (c) Reflectance spectra for control experiment demonstrating minimal change in fringe position when probe-functionalized PSi is exposed to DI water instead of DNase. (d) Reflectance spectra showing redshift of Fabry-Perot fringes after biotin-dsDNA-FAM probe attachment to streptavidin-modified PSi and blueshift of fringes after exposure to DNase and removal of DNA and FAM from the PSi.

To quantitatively relate the observed spectral shifts in Fig. 4(a) to the amount of bound and removed probes, we carried out analytical simulations to model the Fabry-Perot fringe shifts due to biotin-dsDNA-Qdots attachment and DNase cleavage, using a combination of the analytical transfer matrix method and the Bruggeman effective index approximation, as detailed in section 3.6. Figure 4(b) shows that the simulated reflectance spectra are in good agreement with the measured data in Fig. 4(a) when a biotin-dsDNA-Qdot probe fill fraction of *v* = 1.31% is assumed. By considering the footprint of the PSi sample (0.25 cm^2^) and thickness of the PSi film (3.99 mm), as well as the volume of the probes, which includes the volume of the streptavidin-coated Qdots and biotinylated DNA, we then estimated the total number of Qdots bound in the pores to be 0.6 picomoles.

The estimated number of Qdots bound in the pores as determined by the reflectance measurements and simulations was then correlated with the fluorescence data in Fig. 3(b). Using a calibration curve produced with stock Qdot solution, we determined the decrease in fluorescence intensity of the Qdot probe solution after exposure to the PSi sensor during functionalization corresponds to 0.85 picomoles of Qdots in the collected supernatant. Since the probe solution initially contained 3.24 picomoles of Qdots, it can be inferred that the maximum number of Qdots attached in the PSi is 2.34 picomoles. The actual number of Qdots attached in the pores is likely less because the PSi was thoroughly rinsed with DI water after collection of the supernatant and any unbound Qdot probes in the pores would have been rinsed out of the pores but not collected for the fluorescence measurements. Accordingly, we consider the simulations and fluorescence data to be in good qualitative agreement.

There are two key differences between HCPCD and a traditional DNA sensing assay: the direction and magnitude of the spectral change for a detection event. In a traditional sensing assay, DNA hybridization between DNA probe and target molecules causes a spectral redshift. In contrast, for HCPCD, using enzymes to cleave DNA molecules that tether the Qdots to the surface leads to removal of part of the DNA tether and the Qdot from the surface, causing a spectral blueshift. This spectral blueshift – the magnitude of which can be controlled by the choice of the probes size and material – is advantageous for handling biofouling, a major concern for sensors operating in complex media. Since a HCPCD detection event leads to a spectral blueshift, non-specific attachment of proteins or other low-index biomolecule species that result in a biofouling-induced spectral redshift would not cause a false positive sensor response. As long as the sensor surface is stable, as was shown in this work, using a spectral blueshift to signify a detection event has a significant advantage for sensing in saliva, serum, and blood. Further, by taking advantage of the dual amplification achieved from single enzymes cleaving large numbers of probes and the high refractive index of the nanoparticles or Qdots used to label the cleavage probes, it is possible to overcome the sensitivity limitations of traditional biosensing systems.

Since the magnitude of the cleavage-induced spectral change can be increased by the judicious choice of the high-contrast probes, HCPCD leads to a larger signal-to-noise ratio than the standard sensing modality based on detecting low-index biological material added to the system in the process of DNA hybridization. In prior PSi sensor studies, micromolar level target DNA concentrations were required to cause a similar magnitude fringe shift as the DNase cleavage of the 135 nM dsDNA-Qdots reported in this work (Fig. 4(a)) [29], [30]. While DNase is non-selective in cleaving DNA and thus cannot be used for selective detection of DNA sequences unless a more sophisticated detection scheme such as three probe hybridization[31]–[33] is utilized, the work reported here lays an important foundation for HCPCD that could be performed with CRISPR-Cas selective cleavage of nucleic acids using on-chip optical biosensors or other cleavage assays [3], [34], [35] resulting in a highly specific nucleic acid test. The peak shift observed after 2 hours of DNase cleavage at a 12.5 nM concentration is 20 times the resolution of the spectrometer used for the reflectance measurements, showing that there is a path to highly sensitive and fast, real-time readout of assays that result in a cleavage event. Additional optimization may be achieved by decreasing the proportion of ds-DNA linkers to Qdots in order to increase the average refractive index change per occupied streptavidin binding site on the porous silicon surface.

## 5. Conclusion

This work demonstrates the practical potential of high contrast probe cleavage detection as a new sensing modality. PSi interferometers were functionalized with biotin-dsDNA-Qdot probes that served as the high contrast species to be detected when target solution, in this case 12.5 nM DNase, was exposed to the sensor. The magnitude of the measured blueshift after DNA cleavage was much larger than the magnitude of the redshift that would have resulted from hybridization of a nM concentration of DNA in traditional PSi sensing assays. Moreover, we showed that larger and higher refractive index inorganic quantum dots serve as more effective probe labels than smaller and lower refractive index organic FAM dye. HCPCD has the potential to enable new applications for optical biosensors by providing a sensing modality that is robust to noise from surface fouling or complex patient sample media, generates large signal amplification via the use of probes with high-contrast nanoparticles and multiplicative enzymatic activity, and takes advantage of the high target specificity provided by enzymatic assays. This approach can also be used in cleavage based bioassays such as Cas12a and Cas13 CRISPR viral diagnostics, as well as toehold-switch and DNA-zyme biomolecular sensors.

## Acknowledgements

The authors gratefully acknowledge Dmitry Koktysh for his technical assistance with fluorescence measurements and Tengfei Cao for his technical assistance with SEM measurements that were carried out in the Vanderbilt Institute of Nanoscale Science and Engineering. The authors also thank Paul Laibinis for providing access to the ATR-FTIR.

